## Supplementary material for "Tau protein aggregation associated with SARS-CoV-2 main protease"

**Table of contents**

Figure S1. SDS PAGE of 2N4R tau purification.

Figure S2. Characterization of 2N4R tau.

Figure S3. Effect of SARS-CoV-2 3CL$^{pro}$ and tau doses on 2N4R tau aggregation.

Figure S4. Effect of 3CL$^{pro}$ inactivation on tau metabolization.

Figure S5. Mass spectrometry analysis of tau metabolites (GDTPSLEDEAAGHVTQAR).

Figure S6. Mass spectrometry analysis of tau metabolites (SPQLATLADEVSASLAK).

Figure S7. Preferred cleavage sequence pattern of SARS-CoV-2 3CL$^{pro}$ and possible cleavage sites in the tau sequence.

Table S1. Tryptic peptides of 2N4R tau.

Table S2. Tryptic peptides of 2N4R tau detected in peak I.

Table S3. Tryptic peptides of 2N4R tau detected in peak region II.

Table S4. Tryptic peptides of 2N4R tau detected in peak region III.

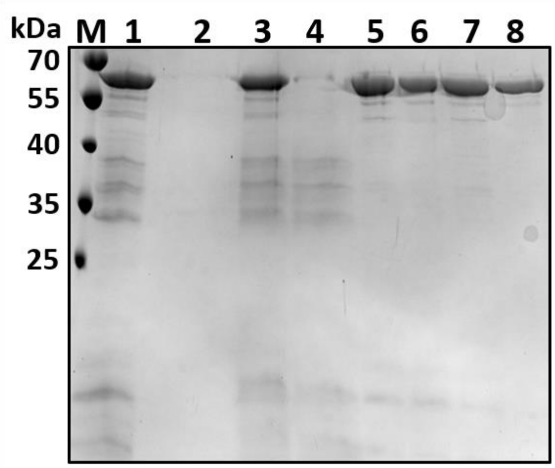


**Supplementary figure S1. SDS PAGE of 2N4R tau purification**. M: Protein marker; 1: Supernatant after cell disruption; 2: Supernatant after ammonium sulfate precipitation; 3: Pellet after ammonium sulfate precipitation solved in buffer 2; 4: SN after centrifugation pellet solved in buffer 2; 5: Pellet solved in dd$H_2O$ and 2 mM TCEP; 6: Pellet was solved in buffer 3; 7: SN after centrifugation pellet solved in buffer 3; 8: Sample after dialysis against buffer 4.

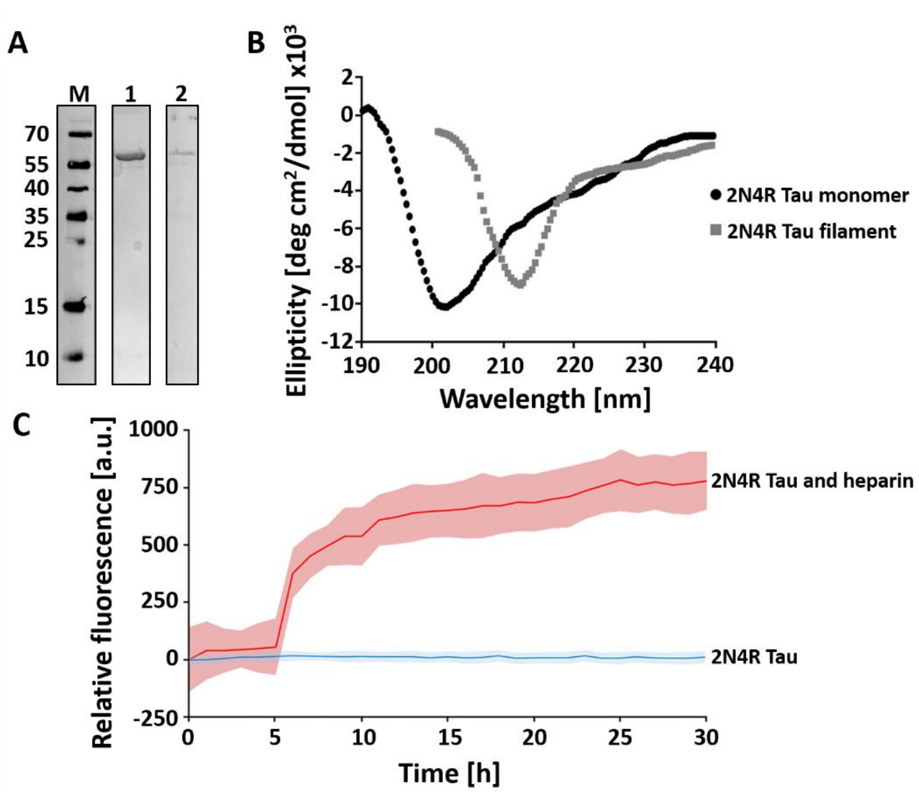


**Supplementary figure S2. Characterization of 2N4R tau. A:** SDS PAGE 15 % of 2N4R tau. M: protein marker, 1: pure 2N4R tau after precipitation purification, 2: western blot with CF633 labeled $tau_{13}$. **B:** CD spectrum of 2N4R tau monomer and filament. **C:** ThT assay of 2N4R tau (10 µM), aggregation was induced with 2.5 µM heparin. Data shown are the mean ± SD from three independent measurements (*n*=3).

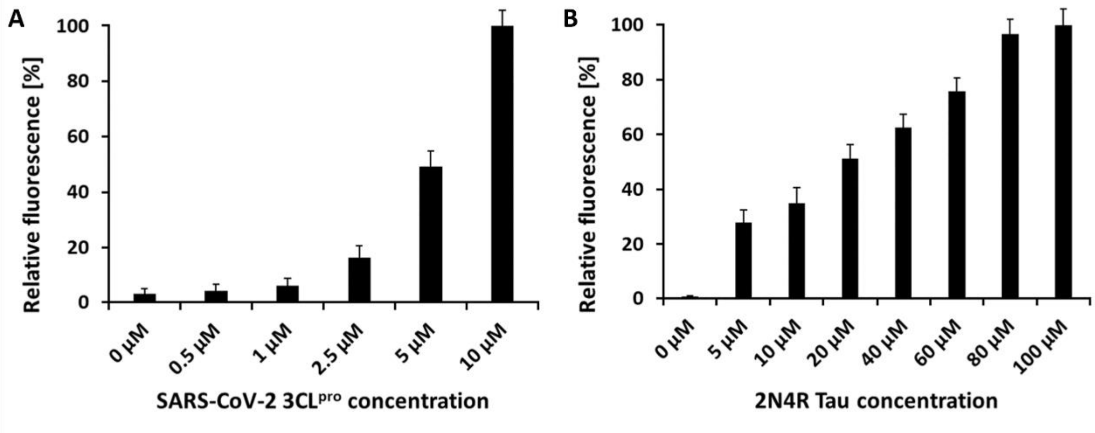


**Supplementary figure S3. Effect of SARS-CoV-2 3CL$^{pro}$ and tau doses on 2N4R tau aggregation.** The endpoint of the relative fluorescence during a ThT assay is shown. **A:** Effect of different 3CL$^{pro}$ concentration (0, 0.5, 1, 2.5, 5 and 10 µM) on tau aggregation (Tau concentration was 10 µM). **B:** Effect of different tau concentration (0, 5, 10, 20, 40, 60, 80 and 100 µM) on tau aggregation. (3CL$^{pro}$ concentration was 10 µM).

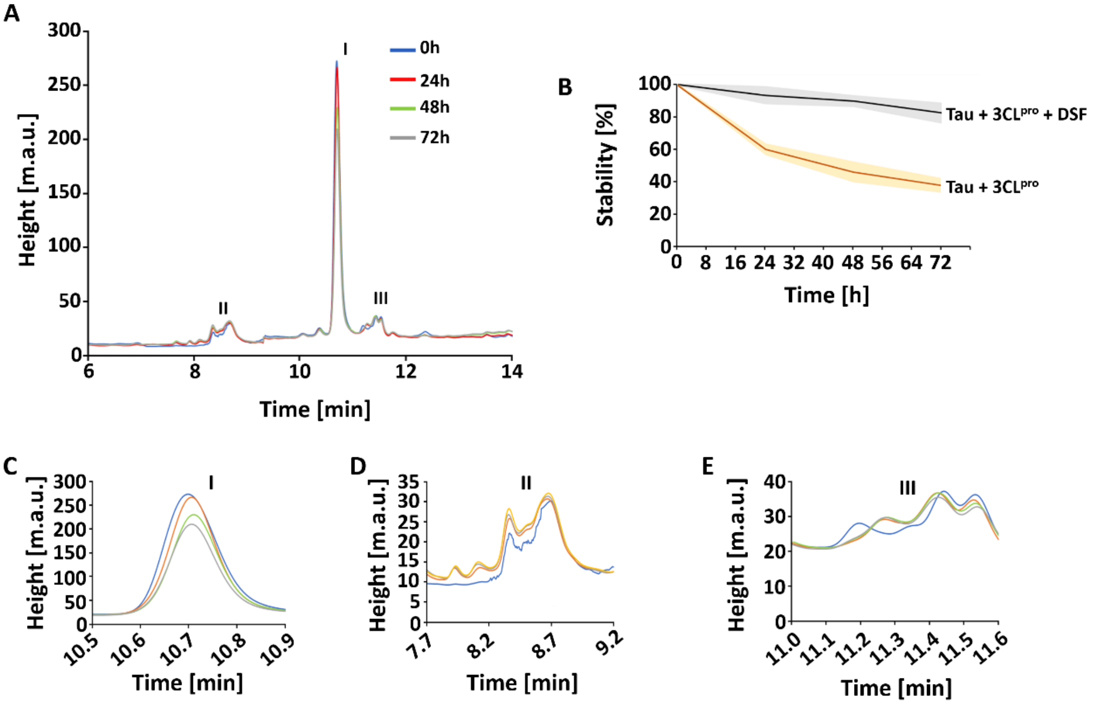


**Supplementary figure S4. Effect of 3CL$^{pro}$ inactivation on tau metabolization.** The protease was inactivated by 10 µM DSF. **A:** Analytical HPLC analysis of 2N4R tau incubated with inactivated SARS-CoV-2 3CL$^{pro}$ for 0, 24, 48 and 72h. The corresponding chromatogram regions of the tau monomer and related metabolites generated by active 3CL$^{pro}$ are highlighted (I-III). **B:** Stability of 2N4R tau monomer after treatment with inactivated 3CL$^{pro}$ over 72h. The tau monomer amount remains at 95 %, compared with a control were tau was treated with active 3CL$^{pro}$. In the control experiment the tau monomer amount reduce to around 40 %. **C:** Chromatogram of peak I (Tau) shown enlarged, **D:** Chromatogram of peak region II shown and **E:** Chromatogram of peak region III shown enlarged. Data shown are the mean ± SD from three independent measurements (*n*=3).

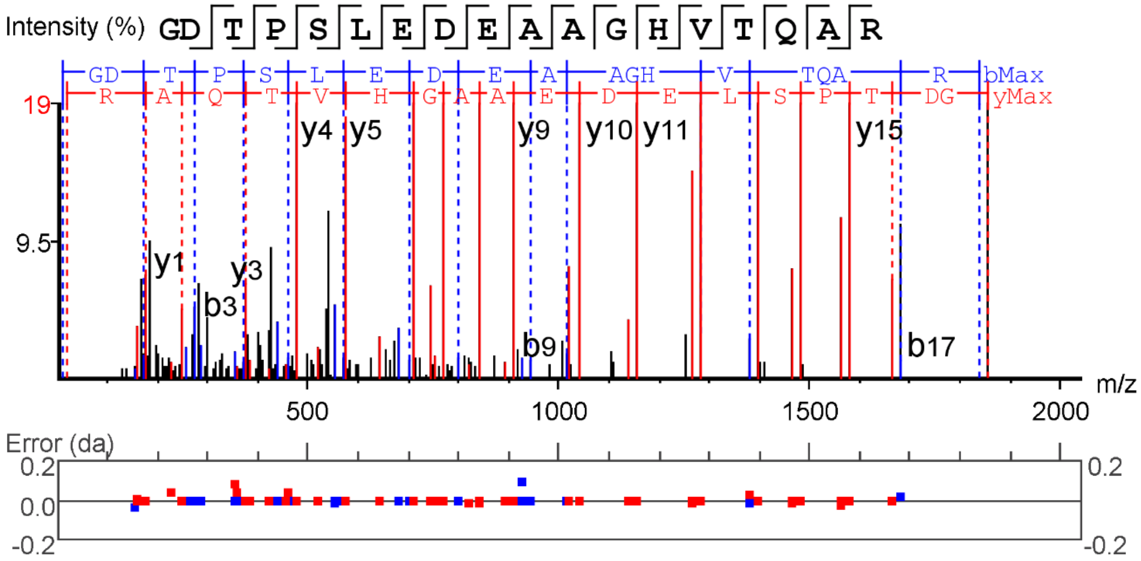


**Supplementary figure S5. Mass spectrometry analysis of tau metabolites.** Example of a MS spectrum corresponding to a tryptic peptide derived from 2N4R tau (GDTPSLEDEAAGHVTQAR), b- and y-ions are labelled.

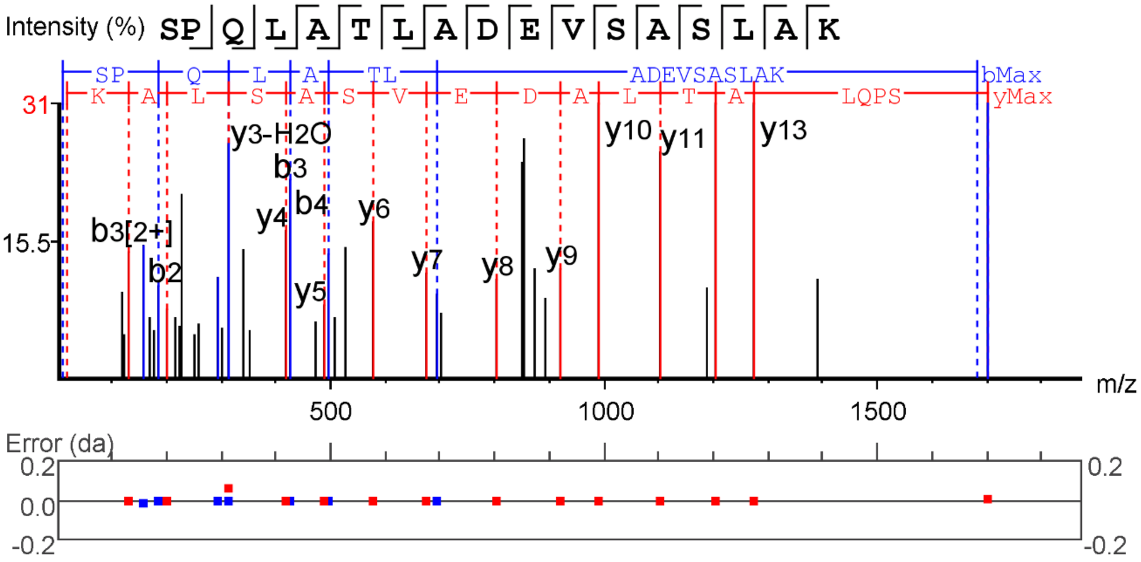


**Supplementary figure S6. Mass spectrometry analysis of tau metabolites.** Example of a MS spectrum corresponding to a tryptic peptide derived from 2N4R tau (SPQLATLADEVSASLAK), b- and y-ions are labelled.

| Substrate binding site SARS-CoV-2 3CL$^{pro}$ | P4 | P3 | P2 | P1 | P1` | P2` | P3` | P4´ | Merops[1] |
|---|---|---|---|---|---|---|---|---|---|
| | A<br>V<br>P<br>T | T<br>K<br>R<br>V<br>M | L<br>I<br>F<br>V | Q | S<br>A<br>G<br>N | A<br>G<br>S<br>I<br>K<br>L<br>N<br>V | E<br>N<br>A<br>D<br>F<br>C<br>M<br>Q<br>V | A<br>S<br>K<br>L<br>N<br>R<br>V | SARS-CoV2 3CL$^{pro}$ |
| Tau$_{6/7}$ | $E_3$ | $P_4$ . | $R_5$ | $Q_6$ * | $E_7$ : | $F_8$ : | $E_9$ * | $V_{10}$ * | No viral protease cleavage site |
| Tau$_{26/27}$ | $R_{23}$ | $K_{24}$ * | $D_{25}$ | $Q_{26}$ * | $G_{27}$ * | $G_{28}$ * | $Y_{29}$ : | $T_{30}$ : | C49.001 |
| Tau$_{33/34}$ | $T_{30}$ * | $M_{31}$ * | $H_{32}$ . | $Q_{33}$ * | $D_{34}$ : | $Q_{35}$ : | $E_{36}$ * | $G_{37}$ . | No viral protease cleavage site |
| Tau$_{35/36}$ | $H_{32}$ | $Q_{33}$ : | $D_{34}$ | $Q_{35}$ * | $E_{36}$ : | $G_{37}$ * | $D_{38}$ * | $T_{39}$ : | No viral protease cleavage site |
| Tau$_{49/50}$ | $S_{46}$ : | $P_{47}$ . | $L_{48}$ * | $Q_{49}$ * | $T_{50}$ : | $P_{51}$ . | $T_{52}$ : | $E_{53}$ : | No viral protease cleavage site |
| Tau$_{88/89}$ | $P_{85}$ * | $G_{86}$ | $K_{87}$ | $Q_{88}$ * | $A_{89}$ * | $A_{90}$ * | $A_{91}$ * | $Q_{92}$ : | C30.001 |
| Tau$_{92/93}$ | $A_{89}$ * | $A_{90}$ : | $A_{91}$ . | $Q_{92}$ * | $P_{93}$ . | $H_{94}$ : | $T_{95}$ : | $E_{96}$ : | No viral protease cleavage site |
| Tau$_{165/166}$ | $Q_{162}$ | $K_{163}$ * | $G_{164}$ | $Q_{165}$ * | $A_{166}$ * | $N_{167}$ * | $A_{168}$ * | $T_{169}$ : | C30.007 |
| Tau$_{244/245}$ | $S_{241}$ : | $R_{242}$ * | $L_{243}$ * | $Q_{244}$ * | $T_{245}$ : | $A_{246}$ * | $P_{247}$ . | $V_{248}$ * | No viral protease cleavage site |
| Tau$_{288/289}$ | $S_{285}$ : | $N_{286}$ * | $V_{287}$ * | $Q_{288}$ * | $S_{289}$ * | $K_{290}$ * | $C_{291}$ * | $G_{292}$ . | C30.001; C30.003; C30.005; C30.007 |
| Tau$_{351/352}$ | $D_{348}$ | $R_{349}$ * | $V_{350}$ * | $Q_{351}$ * | $S_{352}$ * | $K_{353}$ * | $I_{354}$ : | $G_{355}$ . | C30.001; C30.003; C30.005; C30.007 |

**Supplementary figure S7. Preferred cleavage sequence pattern of SARS-CoV-2 3CL$^{pro}$ and possible cleavage sites in the tau sequence**. 3CL$^{pro}$ substrate binding site and preferred amino acids are shown and sequence homology of possible cleavage sites. The identical amino acid pattern was checked in the Merops database (Rawlings et al., 2018) if there exist identities to known viral protease cleavage sequences. The following virus proteases were identified: C49.001 (Strawberry mottle virus 3C-like peptidase); C30.001 (Coronavirus picornain 3C-like peptidase-1); C30.003 (Human coronavirus 229E main peptidase); C30.005 (SARS coronavirus picornain 3C-like peptidase) and C30.007 (Coronavirus COVID-19 3C-like peptidase).

**Table S1.** Tryptic peptides of 2N4R tau.

| | 2N4R tau | m/z | ppm | length | Mass | Feature | Accession |
|---|---|---|---|---|---|---|---|
| 1 | AEPRQEFEVMEDHAGTYGLGDR | 8363807 | 9 | 22 | 25061182 | 8 | Human_TAU |
| 2 | GIGDTPSLEDEAAGHVTQAR | 10124886 | 11 | 20 | 20229606 | 7 | Human_TAU |
| 3 | HVPGGGSVQIVY | 6068225 | 5 | 12 | 12116299 | 7 | Human_TAU |
| 4 | GDTPSLEDEAAGHVTQAR | 9274343 | -6 | 18 | 18528551 | 6 | Human_TAU |
| 5 | TPSLEDEAAGHVTQAR | 8414106 | 0 | 16 | 16808066 | 6 | Human_TAU |
| 6 | AGLKESPLQTPTEDGSEEPGSETSDAK | 13806379 | -2 | 27 | 27592620 | 5 | Human_TAU |
| 7 | TAPVPMPDLK | 5347914 | -3 | 10 | 10675685 | 5 | Human_TAU |
| 8 | TDAGLKESPLQTPTEDGSEEPGSETSDAK | 14886765 | 6 | 29 | 29753367 | 4 | Human_TAU |
| 9 | AEPRQEFEVMEDHAGTYGLGDRK | 8790784 | 1 | 23 | 26342131 | 4 | Human_TAU |
| 10 | STPTAEDVTAPLVDEGAPGKQ | 10420134 | 3 | 21 | 20820117 | 4 | Human_TAU |
| 11 | SLDNITHVPGGGNK | 7048621 | -7 | 14 | 14077106 | 4 | Human_TAU |
| 12 | SPQLATLADEVSASLAKQGL | 10000389 | 0 | 20 | 19980632 | 4 | Human_TAU |
| 13 | AEPRQEFEVMEDHAGTY | 10049391 | 2 | 17 | 20078632 | 4 | Human_TAU |
| 14 | PSLPTPPTREPK | 4405823 | 4 | 12 | 13187245 | 4 | Human_TAU |
| 15 | PRQEFEVMEDHAGTYGLGDR | 7696855 | -16 | 20 | 23060386 | 3 | Human_TAU |
| 16 | TPPSSGEPPKSGDRSGYSSPGSPGTPGSR | 9294379 | 5 | 29 | 27852903 | 3 | Human_TAU |
| 17 | TDHGAEIVYKSPVVSGD | 8874371 | 9 | 17 | 17728580 | 3 | Human_TAU |
| 18 | SKIGSLDNITHVPGGGNK | 8974785 | -4 | 18 | 17929431 | 3 | Human_TAU |
| 19 | GSLGNIHHKPGGGQVEVK | 9074884 | 15 | 18 | 18129595 | 3 | Human_TAU |
| 20 | IGSLDNITHVPGGGN | 7258679 | 1 | 15 | 14497212 | 3 | Human_TAU |
| 21 | QTAPVPMPDLK | 5988204 | -7 | 11 | 11956271 | 3 | Human_TAU |
| 22 | QIVYKPVDLSK | 6453762 | -9 | 11 | 12887390 | 3 | Human_TAU |
| 23 | IVYKPVDLSK | 5813475 | 0 | 10 | 11606804 | 3 | Human_TAU |
| 24 | PVPMPDLK | 4487482 | -21 | 8 | 8954837 | 3 | Human_TAU |
| 25 | SLPTPPTREPK | 6118436 | 8 | 11 | 12216716 | 3 | Human_TAU |
| 26 | QEFEVMEDHAGTY | 7783214 | -9 | 13 | 15546296 | 3 | Human_TAU |
| 27 | PTPPTREPK | 3415256 | -7 | 9 | 10215556 | 3 | Human_TAU |

**Table S2.** Tryptic peptides of 2N4R tau detected in peak I.

| | Peak I | m/z | ppm | length | Mass | Feature | Accession |
|---|---|---|---|---|---|---|---|
| 1 | SPQLATLADEVSASLAK | 8509570 | 2 | 17 | 16998992 | 8 | Human_TAU |
| 2 | GDTPSLEDEAAGHVTQAR | 9274293 | -59 | 18 | 18528551 | 5 | Human_TAU |
| 3 | PTPPTREPK | 5117853 | 4 | 9 | 10215556 | 5 | Human_TAU |
| 4 | AGLKESPLQTPTEDGSEEPGSETSDAK | 9207599 | -14 | 27 | 27592620 | 4 | Human_TAU |
| 5 | PTAEDVTAPLVDEGAPGK | 8839431 | -10 | 18 | 17658733 | 4 | Human_TAU |
| 6 | TPSLEDEAAGHVTQAR | 8414100 | -7 | 16 | 16808066 | 4 | Human_TAU |
| 7 | GIGDTPSLEDEAAGHVTQAR | 10124829 | -46 | 20 | 20229606 | 4 | Human_TAU |
| 8 | SPQLATLADEVSASLAKQGL | 10000392 | 3 | 20 | 19980632 | 4 | Human_TAU |
| 9 | AEPRQEFEVMEDHAGTYGLGDR | 8363809 | 10 | 22 | 25061182 | 3 | Human_TAU |
| 10 | DTPSLEDEAAGHVTQAR | 8989072 | -177 | 17 | 17958336 | 3 | Human_TAU |
| 11 | SLDNITHVPGGGNK | 7048618 | -11 | 14 | 14077106 | 3 | Human_TAU |
| 12 | VQIVYKPVDLSK | 4636093 | -11 | 12 | 13878075 | 3 | Human_TAU |
| 13 | PSLPTPPTREPK | 4405822 | 2 | 12 | 13187245 | 3 | Human_TAU |
| 14 | IVYKPVDLSK | 5813471 | -7 | 10 | 11606804 | 3 | Human_TAU |
| 15 | SLPTPPTREPK | 6118428 | -5 | 11 | 12216716 | 3 | Human_TAU |

**Table S3.** Tryptic peptides of 2N4R tau detected in peak region II.

| | **Peak region II** | **m/z** | **ppm** | **length** | **Mass** | **Feature** | **Accession** |
|---|---|---|---|---|---|---|---|
| 1 | GDTPSLEDEAAGHVTQAR | 9274330 | -20 | 18 | 18528551 | 6 | Human_TAU |
| 2 | GIGDTPSLEDEAAGHVTQAR | 10124861 | -14 | 20 | 20229606 | 4 | Human_TAU |
| 3 | STPTAEDVTAPLVDEGAPGKQ | 10420308 | 169 | 21 | 20820117 | 4 | Human_TAU |
| 4 | SPQLATLADEVSASLAK | 8509573 | 5 | 17 | 16998992 | 4 | Human_TAU |
| 5 | PTAEDVTAPLVDEGAPGK | 8839406 | -38 | 18 | 17658733 | 3 | Human_TAU |
| 6 | LATLADEVSASLAK | 6948846 | -8 | 14 | 13877559 | 3 | Human_TAU |
| 7 | SLDNITHVPGGGNK | 7048621 | -7 | 14 | 14077106 | 3 | Human_TAU |
| 8 | HVPGGGSVQIVY | 6068227 | 8 | 12 | 12116299 | 3 | Human_TAU |
| 9 | GNIHHKPGGGQVEVK | 5196147 | 2 | 15 | 15558219 | 3 | Human_TAU |
| 10 | GSVQIVYKPVDLSK | 7669375 | -3 | 14 | 15318610 | 3 | Human_TAU |
| 11 | PTPPTREPK | 5117839 | -22 | 9 | 10215556 | 3 | Human_TAU |

**Table S4.** Tryptic peptides of 2N4R tau detected in peak region III.

| | **Peak region III** | **m/z** | **ppm** | **length** | **Mass** | **Feature** | **Accession** |
|---|---|---|---|---|---|---|---|
| 1 | SPQLATLADEVSASLAK | 5676406 | 5 | 17 | 16998992 | 15 | Human_TAU |
| 2 | SLGNIHHKPGGGQVEVK | 8789772 | 10 | 17 | 17559380 | 4 | Human_TAU |
| 3 | SLDNITHVPGGGNK | 7048634 | 12 | 14 | 14077106 | 4 | Human_TAU |
| 4 | SLDNITHVPGGGNKK | 5129423 | -3 | 15 | 15358055 | 4 | Human_TAU |
| 5 | GDTPSLEDEAAGHVTQAR | 9274317 | -34 | 18 | 18528551 | 3 | Human_TAU |
| 6 | SNVSSTGSIDMVDSPQLATLADEVSASLAK | 9984936 | 15 | 30 | 29924546 | 3 | Human_TAU |
| 7 | GIGDTPSLEDEAAGHVTQAR | 10124868 | -8 | 20 | 20229606 | 3 | Human_TAU |
| 8 | SPQLATLADEVSASLAKQGL | 10000397 | 8 | 20 | 19980632 | 3 | Human_TAU |
| 9 | PGGGSVQIVYKPVDLSK | 8724852 | -4 | 17 | 17429567 | 3 | Human_TAU |
| 10 | DTPSLEDEAAGHVTQAR | 8989229 | -13 | 17 | 17958336 | 3 | Human_TAU |
| 11 | IGSLDNITHVPGGGN | 7258684 | 7 | 15 | 14497212 | 3 | Human_TAU |
| 12 | GGGSVQIVYKPVDLSK | 8239599 | 9 | 16 | 16459038 | 3 | Human_TAU |
| 13 | QIVYKPVDLSK | 6453778 | 16 | 11 | 12887390 | 3 | Human_TAU |
| 14 | VYKPVDLSK | 5248058 | 6 | 9 | 10475964 | 3 | Human_TAU |
| 15 | SLPTPPTREPK | 6118428 | -5 | 11 | 12216716 | 3 | Human_TAU |
| 16 | PTPPTREPK | 5117839 | -24 | 9 | 10215556 | 3 | Human_TAU |
